# Unexpected Detection of Highly Pathogenic Avian Influenza (HPAI) H5N1 virus in bovine semen from a bull used for natural breeding on an affected dairy farm

**DOI:** 10.1101/2025.10.16.682947

**Authors:** Ailam Lim, Keith Poulsen, Leonardo C. Caserta, Lizheng Guan, Eryn Opgenorth, Maxwell P. Beal, Amie J. Eisfeld, Yoshihiro Kawaoka, Diego G. Diel

**Affiliations:** University of Wisconsin-Madison Wisconsin Veterinary Diagnostic Laboratory, Madison, Wisconsin, USA; University of Wisconsin-Madison Dept. of Pathobiological Sciences, Madison, Wisconsin, USA; University of Wisconsin-Madison Dept. of Medical Sciences, Madison, Wisconsin, USA; Cornell University Animal Health Diagnostic Center, Department of Population Medicine and Diagnostic Sciences, College of Veterinary Medicine, Ithaca, New York, USA; University of Wisconsin-Madison, Dept. of Pathobiological Sciences and Influenza Research Institute, Madison, Wisconsin, USA; Mill Creek Veterinary Services, Visalia, California, USA; Dakota Dairy Health, Brookings, South Dakota, USA; University of Tokyo Department of Virology, Institute of Medical Science, Tokyo, Japan; University of Tokyo Pandemic Preparedness, Infection and Advanced Research Center (UTOPIA), Tokyo, Japan; Japan Institute for Health Security, National Institute of Global Health and Medicine, Tokyo, Japan

**Keywords:** Highly Pathogenic Avian Influenza, H5N1, Biosecurity

## Abstract

Since March 2024, HPAI H5N1 virus has infected dairy cattle in the U.S., prompting concern about novel transmission routes. During an outbreak in California, HPAI H5N1 RNA was detected in an asymptomatic bull’s semen. Although infectious virus was not isolated, questions remain about semen-associated transmission risks and biosecurity practices.

## Main Text

Since March 2024, detection of clade 2.3.4.4b highly pathogenic avian influenza (HPAI) H5N1 in U.S. dairy cattle has raised concerns about the virus’s ability for cross-species transmission, adaptation to mammals, and novel transmission routes, including milk (1,2). The knowledge that viruses are infectious in bovine semen and the finding of HPAI in turkey semen, has prompted questions about the potential role of HPAI transmission in bovine semen (3,4). The shedding of HPAI H5N1 in bovine semen could result in silent viral spread within herds and across geographic regions via artificial insemination (AI). While there are reports of clinical HPAI disease in female calves and pregnant animals, reports of diseased bulls in dairies or beef cattle are lacking. While many questions about the pathophysiology of HPAI H5N1 in U.S. dairy herds remain unanswered, movement of lactating cows is a recognized risk factor for interstate disease spread. This diagnostic study sought evidence of HPAI H5N1 shed via semen in natural breeding bulls on an HPAI H5N1-affected dairy farm in California.

The HPAI H5N1 genotype B3.13 outbreak in California began in August 2024, likely resulting from the interstate movement of infected cows, which led to the rapid spread of the virus within the state. In October 2024, infection in a 4,500-head Holstein dairy was detected by reverse transcription-polymerase chain reaction (RT-PCR) for H5N1 RNA in bulk tank milk samples. Clinical signs in lactating dairy cows included decreased milk production, mastitis, lethargy, dehydration, anorexia, and pyrexia (>104.0°F). The herd experienced a 60% morbidity rate over three weeks. About four weeks later, the herd veterinarian collected diagnostic samples from three three-year-old Holstein bulls, including deep nasal swabs, preputial scrapings, pre-ejaculate seminal fluid, semen and serum samples. These were sent to the Wisconsin Veterinary Diagnostic Laboratory for testing. Subjectively, the semen appeared to have low sperm concentration and volume, although total sperm count was not measured. The poor semen quality was likely due to timing and suboptimal sampling conditions, as the bulls had been co-mingled with cows the same day for breeding.

Initially, the samples were examined for influenza A virus (IAV) using various methods (Table 1 and Supplementary File 1). Deep nasal swabs, preputial scrapes, pre-ejaculate seminal fluids, and semen were tested using multiple IAV Matrix RT-PCR assays (5). The IAV RNA was detected in the semen from Bull 1 (147816-25) by three different PCR assays, but Bulls 2 and 3 were tested negative. No detection was reported in any other samples. The IAV RNA detection in Bull 1 was confirmed with an additional RNA extraction to rule out laboratory contamination. The IAV strain was further identified as HPAI using the H5N1 2.3.4.4b lineage subtyping RT-PCR assay.

**Table 1:**
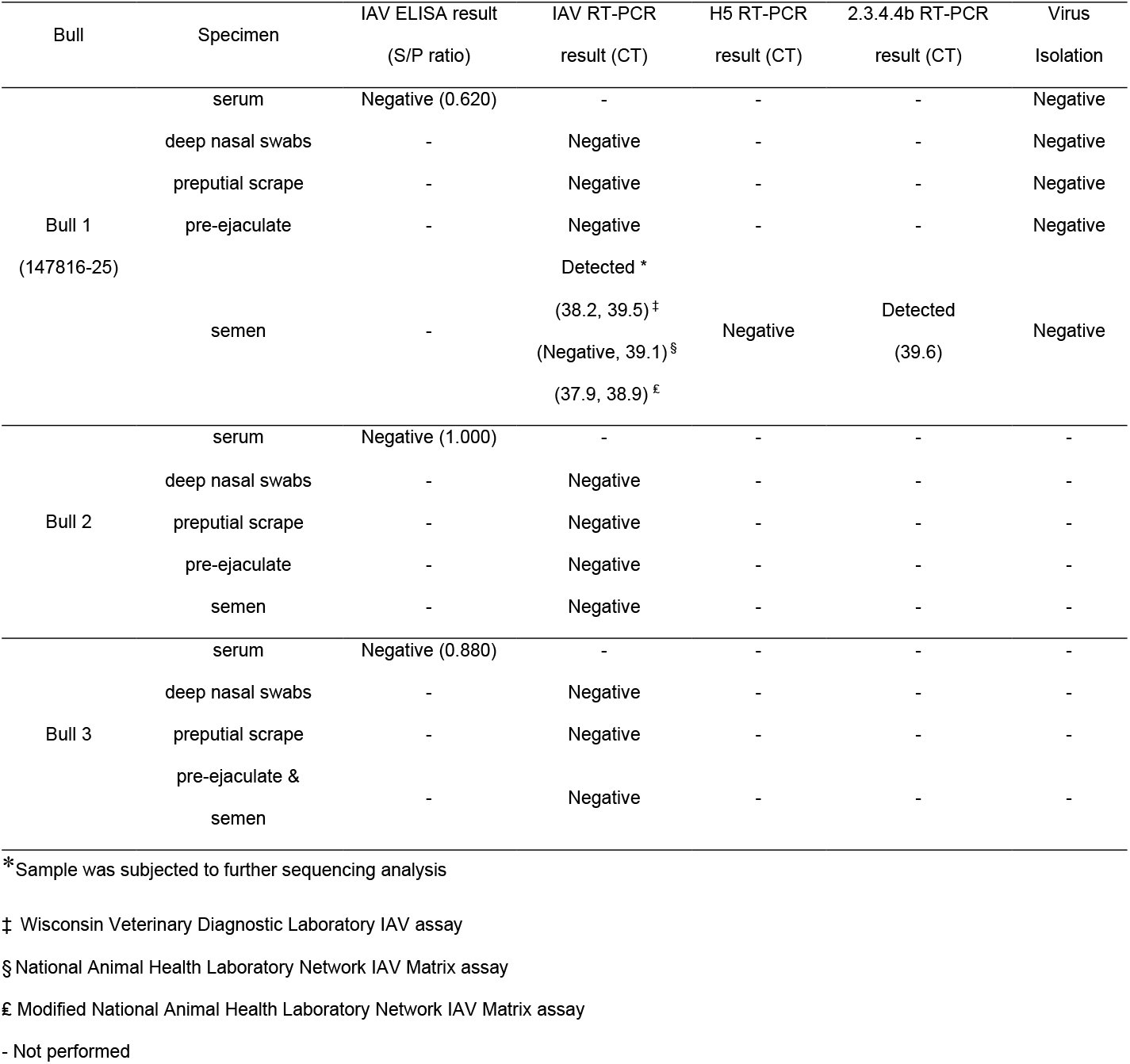
Influenza A virus testing results for the samples collected from the three bulls in an H5N1 infected farm in Central Valley of California.

Targeted IAV sequencing at the Cornell Animal Health Diagnostic Center confirmed the presence and yielded a partial H5N1 genome (Supplementary Table S1, GISAID accession number EPI_ISL_20206713). Attempts to assemble the full genome were unsuccessful due to the low viral load in the semen. Phylogenetic analysis of the partial concatenated genome sequences indicated that the viral RNA in Bull 1 (147816-25) clustered within the B3.13 genotype and was closely related to a sample collected from a Californian dairy farm worker during the same period (Figure 1) (6).

**Figure 1.**
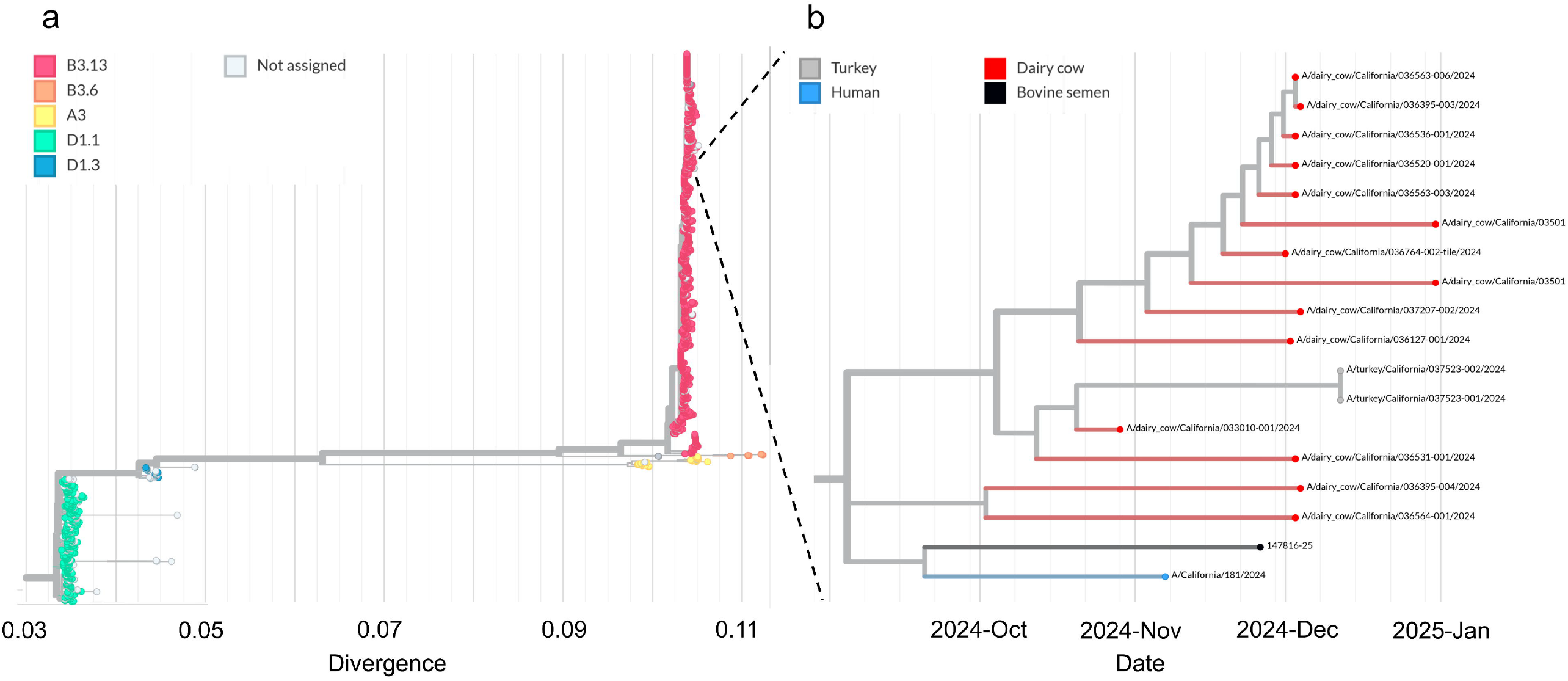
Phylogenetic analysis of partial concatenated genome sequences confirming detection of highly pathogenic avian influenza A(H5N1) virus genotype B3.13 from bovine semen. A) Tree showing broader phylogeny of H5N1 virus genotypes; B) timescale tree showing closer examination of the virus from the Bull 1 semen (sample no. 147816-25) and closely related virus sequences.

All Bull 1 samples were processed for virus isolation at the University of Wisconsin— Madison Influenza Research Institute to attempt recovery of an isolate. Serially diluted samples inoculated into 10-day-old specific-pathogen-free embryonated chicken eggs or Madin–Darby canine kidney cells showed no embryo death or cytopathic effect, respectively. All samples were confirmed negative by hemagglutination assay.

The sera from all bulls tested negative by antigen-based influenza A ELISA (7). However, the S/P ratio for Bull 1 was 0.620, which is close to the validated 0.5 assay cutoff, potentially indicating seroconversion.

Because of the limited semen volume for analysis, further confirmation testing at the national reference laboratory was not performed. Additional samples were requested several months later for convalescent testing, but Bull 1 had been culled from the herd. The importance of identifying HPAI H5N1 in bovine semen remains uncertain, as the virus could have been actively shed in semen or the ejaculate could have been contaminated by the environment. Although detecting RNA does not confirm the presence of infectious viruses, this finding warrants further investigation into whether HPAI H5N1 can be shed in semen and raises questions about farm biosecurity amid the ongoing outbreak. Good biosecurity measures are essential to preventing infections and, if infection occurs, slowing disease spread on the farm.

If these findings were to be repeated and confirmed, it would have significant implications for natural breeding and biosecurity for AI collection centers. Further research and risk assessments are needed to determine whether naturally infected HPAI H5N1 bulls shed virus in semen, and, if so, evaluate the risk of disease spread on dairy farms and with AI programs.

