## Supplementary File 1 for "Unexpected Detection of Highly Pathogenic Avian Influenza (HPAI) H5N1 virus in bovine semen from a bull used for natural breeding on an affected dairy farm"

### Materials and Methods

**Sample collection and preparation.** Due to movement restrictions in effect during the HPAI outbreak in cattle, sampling of the bulls was conducted on-farm. Given the lack of facilities to handle the 2500-3000 lb full-grown, commercial, Holstein bulls, a mobile chute and temporary fencing were set up to corral the bulls. Once in the chute, the bulls' size posed a significant challenge to proper restraint. Two of the five bulls were excluded from sampling due to safety concerns for the bulls, the workers and the attending veterinarian.

Serum was collected in no-additive red-top tubes from the remaining three bulls. Deep nasal swabs and preputial scrapings from all three bulls were collected into 3 ml brain heart infusion (BHI) media. Pre-ejaculate seminal fluid and semen were collected separately from Bull 1 and 2, while a mixture of pre-ejaculate and semen from Bull 3, using electroejaculation with a non-aseptic technique (excess hair was clipped, and the area was not cleaned). It was an onerous task to move the untamed bulls through the chutes for sample collection, the entire process took a veterinarian and 5 dairy staff about 3 hours to complete. All animals were returned to their respective pens without complication.

Samples were shipped on ice packs to Wisconsin Veterinary Diagnostic Laboratory. Upon arrival, serum was separated from clots and store at 4 °C until testing. Preputial scrape was pelleted by centrifugation at 12,000 xg, supernatant was transferred into a new tube and pellet was resuspend in 500 µL of 1x phosphate buffered saline (PBS) for storage at -80 °C. Pre-ejaculate seminal fluid and semen were storage at -80 °C until tested.

**DNA extraction and RT-PCR assays.** Samples were extracted using the IndiMag Pathogen Kit (Indical BIOSCIENCE, Lepzig, Germany). A 200 µL volume input of media for deep nasal swabs or preputial scrapes were used, while 100 µL of resuspended preputial scrape pellets, pre-ejaculate or semen were diluted 1:2 in 1x PBS for extraction (8); Assay specific Internal Positive Controls were added to the lysis solution. The extraction process was conducted according to the manufacturer's instructions using the KingFisher Flex extractor (Thermo Fisher Scientific, Waltham, MA). Eluted RNA was used for RT-PCR evaluation or stored in at −80 °C.

The RNA extracted was evaluated using three IAV RT-PCR assays. The Wisconsin Veterinary Diagnostic Laboratory in-house IAV assay was conducted as published (8); the National Animal Health Laboratories Network (NAHLN) IAV Matrix RT-PCR assay is conducted per NAHLN protocol NVSL-SOP-0068 (5); while the Modified NAHLN IAV Matrix assay was an approved deviation to substitute the AgPath-ID One-Step RT-PCR Reagents (Thermo Fisher Scientific, Waltham, MA) with the IndiMix JOE master mix (Indical BIOSCIENCE, Lepzig, Germany) in a 20 µl reaction (unpublished). Upon positive detection on the IAV assays, subtyping RT-PCR assays for the H5 subtype and the H5N1 2.3.4.4b lineage were carried out according to protocols in the NVSL-SOP-0068. The NAHLN-approved protocols are available online: https://www.aphis.usda.gov/animal_health/lab_info_services/downloads/ApprovedSOPList.pdf. The NAHLN program office controls the distribution of protocols and the detailed information in the previously listed protocols; these can be requested by emailing.

**Targeted influenza A sequencing and bioinformatic analyses**. The semen sample was diluted 1:2 in PBS, and 200 µl of this dilution was used for extraction with the IndiMag Pathogen Kit (Indical BIOSCIENCE, Lepzig, Germany) on the KingFisher Flex extractor (Thermo Fisher Scientific, Waltham, MA), as well as manually extracted using the QIAamp MinElute Virus Spin Kit (Qiagen Science, Germantown, MD). A tiled-amplicon approach was used to amplify all segments of H5N1 viruses using two primer pools (primer sequences available upon request). Sequencing libraries were prepared using the Native Barcoding Kit EXP-NBD196 and the Ligation Sequencing Kit SQK-LSK109 (Nanopore Technologies, Oxford, United Kingdom), and sequenced on a FLO-MIN106 MinION flow cell (R9.4.1) using the GridION platform (Nanopore Technologies, Oxford, United Kingdom). Quality-filtered and primer-trimmed reads were aligned to a reference genome downloaded from GenBank using Minialign software (v0.4.4; https://github.com/ocxtal/minialign). Consensus sequences were generated using Medaka (v1.4.3) with the medaka_haploid_variant and medaka_consensus programs for polishing (https://github.com/nanoporetech/medaka). The dataset used for analysis consisted of HPAI H5N1 genomes from samples collected between August and December 2024 in North America, downloaded from the GISAID EpiFlu database. Phylogenomic analyses were performed using Nextstrain (9), with a modification to input a maximum-likelihood phylogenetic tree of concatenated genomes inferred using IQ-TREE (10) with an edge-linked partition model and 1,000 bootstrap replicates.

**Viral Isolation.** Specific-pathogen-free (SPF) embryonated chicken eggs (VALO BioMedia, Adel, IA) were incubated at 37 °C with appropriate humidity. Ten-day-old eggs were candled to verify embryo viability prior to inoculation. Madin-Darby canine kidney (MDCK) cells were grown in Eagle's minimal essential medium (MEM) containing 5% newborn calf serum and antibiotics at 37 °C in a humidified atmosphere of 5% $CO_2$. Cells were routinely monitored for mycoplasma contamination.

After thawing, liquid samples (deep nasal swabs, preputial wash, pre-ejaculate, and semen) were used as-is. The thawed preputial cell pellet was resuspended in 0.5 ml of phosphate-buffered saline (PBS). All samples were then serially diluted ($10^{-1}$ through $10^{-5}$) in PBS and used to inoculate SPF eggs ($10^{0}$ through $10^{-2}$ dilutions, 100 µl per egg, 2 eggs per dilution) or MDCK cells in 6-well plates ($10^{0}$ through $10^{-5}$ dilutions, 500 µl per well, 1 well per dilution). For MDCK cells, inoculums were removed after 1 hour of incubation at 37 °C, cells were washed once with virus growth medium (MEM containing 0.3% bovine serum albumin and 0.6 µg/ml of L-(tosylamido-2-phenyl) ethyl chloromethyl ketone (TPCK)-treated trypsin), and then covered with virus growth medium. Eggs were incubated as described above for three days, candled to assess viability, and then killed by incubation at 4 °C overnight. MDCK cells were incubated as described above for five days and observed daily by microscopy to assess cytopathic effects.

**Hemagglutination assays.** Allantoic fluids from eggs or MDCK cell culture supernatants were subjected to hemagglutination (HA) assays according to standard methods. Briefly, allantoic fluids or MDCK supernatants were two-fold serially diluted and mixed with 0.5% turkey red blood cells, followed by assessment for agglutination.

**Bovine influenza A ELISA.** Sera were tested at 1:5 dilution using the IDEXX Al MultiS-Screen Enzyme-Linked lmmunosorbent Assay (IDEXX, Westbrook, ME, USA). The test was performed per manufacturer’s instructions, and interpretation per the NAHLN protocol NVSL-SOP-1255 (7) where samples with S/P ratio $\geq 0.5$ are considered absence of influenza A specific antibodies.

**Biosafety.** Attempted virus isolation from bull samples was carried out in a Biosafety Level 3 (BSL-3) containment laboratory at the Influenza Research Institute at the University of Wisconsin-Madison, which is approved by the Federal Select Agent Program for studies with highly pathogenic avian influenza viruses. The University of Wisconsin-Madison Institutional Contact for Dual Use Research reviewed this manuscript and confirmed that the studies described herein do not meet the criteria of Dual Use Research of Concern (DURC).

**Supplementary Table S1**. Genome coverage of the eight influenza A segments for sample Bull 1 (147816-25) generated using a tiled-amplicon approach on a long-read sequencing technology using MinION flow cell (R9.4.1) on the Oxford Nanopore GridION platform. The sequence generated was deposited in GISAID (accession number EPI_ISL_20206713).

| Segment # | Segment Name | # of reads assembled to reference | Segment % Coverage |
|---|---|---|---|
| 1 | Polymerase basic 2 (PB2) | 35992 | 61 |
| 2 | Polymerase basic 1 (PB1) | 2431 | 19 |
| 3 | Polymerase acidic (PA) | 5972 | 47 |
| 4 | Hemagglutinin (HA) | 6342 | 33 |
| 5 | Nucleocapsid (NP) | 7717 | 47 |
| 6 | Neuraminidase (NA) | 11616 | 72 |
| 7 | Matrix (M) | 1019 | 44 |

| | | | |
|---|---|---|---|
| 8 | Nonstructural (NS) | 10182 | 60 |
